# Coordination of membrane synthesis with cell growth: the *Escherichia coli* phospholipid synthesis enzyme PlsB responds to membrane abundance in a manner consistent with filamentation-mediated inhibition

**DOI:** 10.1101/2025.11.03.685888

**Authors:** Jaïrus Beije, Desi Fierlier, Margot Guurink, Amba Stapert, Adja Zoumaro-Djayoon, Gregory Bokinsky

## Abstract

Cell viability demands tight coordination between growth and the synthesis of new membrane. The mechanisms coordinating cell growth with membrane synthesis in any organism are unclear. In *Escherichia coli,* initiation of phospholipid synthesis by the glycerol-3-phosphate acyltransferase PlsB is regulated by cell growth via an unknown allosteric mechanism. Previous studies have established that PlsB assembles into enzymatically inactive, membrane-bound filaments when overexpressed. We propose that PlsB filamentation regulates PlsB activity and coordinates membrane synthesis with growth. Here, we test our hypothesis by observing the localization of fluorescently labelled PlsB using live-cell fluorescence microscopy. During growth, PlsB localizes as discrete foci, consistent with formation of inactive filaments. Reducing cellular membrane content eliminates PlsB foci and delocalizes PlsB into the cytoplasm. Restoring membrane synthesis causes PlsB foci to reform after a delay. These results are consistent with our hypothesis and suggest a model in which PlsB reversibly assembles into inactive filaments in response to phospholipid abundance. This mechanism establishes a negative feedback loop that controls initiation of phospholipid synthesis by PlsB and effectively coordinates membrane synthesis with growth.

## Introduction

How do cells grow without breaking their membranes? Cells must tightly coordinate cell envelope synthesis with growth to maintain viability. This ability is essential for all cells and yet is not understood in any organism. The membrane synthesis pathway of the model prokaryote *Escherichia coli* is perhaps the best-characterized to date (1). *E. coli* is a Gram-negative bacterial species whose inner membrane is composed of phospholipids while the outer membrane is an asymmetric bilayer featuring a phospholipid inner leaflet and a lipopolysaccharide outer leaflet. The synthesis of membrane phospholipids is initiated in the cytoplasm by the peripheral membrane enzyme PlsB, which transfers fatty acids from acyl thioesters to glycerol-3-phosphate (G3P) to generate lysophosphatidic acid (LPA) (**Fig. 1A**). LPA is further acylated by PlsC to generate phosphatidic acid (PA), the universal precursor of glycerophospholipids (2). Subsequent reactions add headgroups to the G3P backbone, yielding membrane phospholipids phosphatidylglycerol (PG) and phosphatidylethanolamine (PE). While LPA is also generated via PlsX and PlsY, this pathway is not essential in *E. coli,* indicating that most LPA is synthesized by PlsB (3). Synthesized phospholipids partition into the inner membrane and are subsequently transferred to the inner leaflet of the outer membrane by several redundant protein bridges (4).

**Figure 1.**
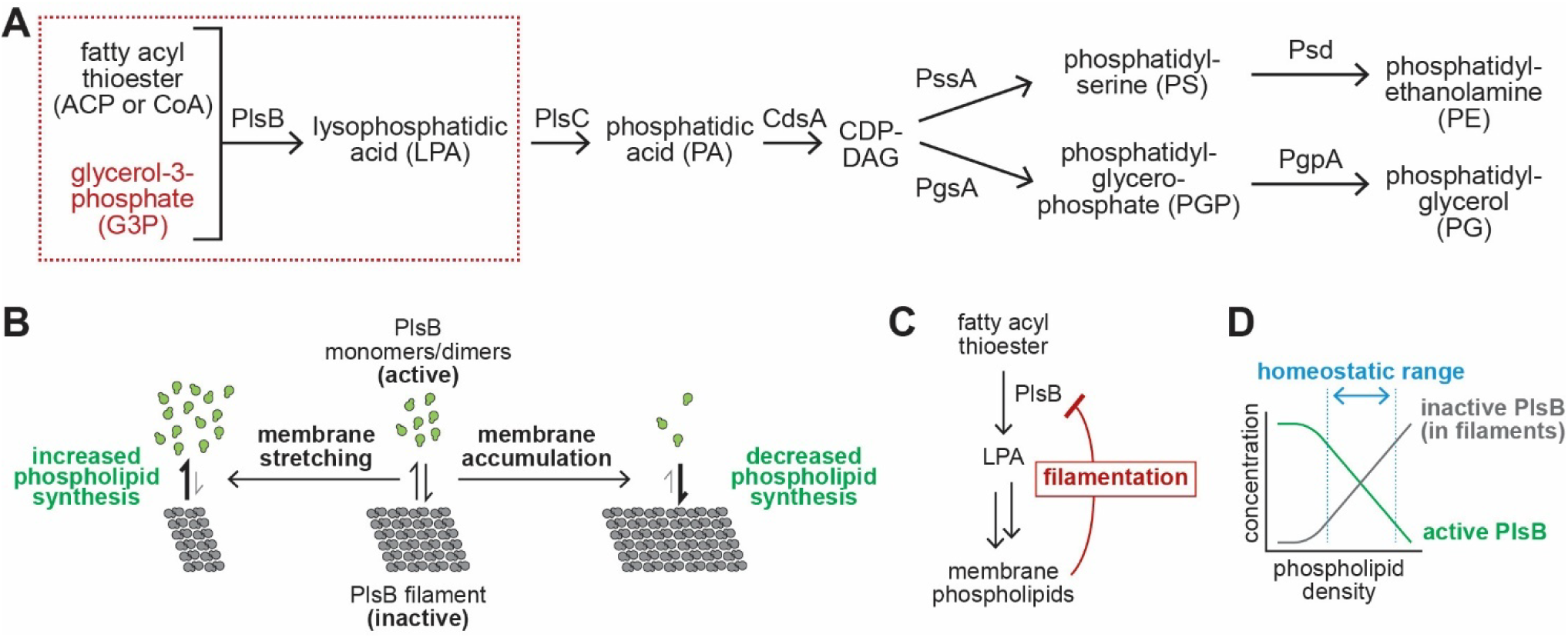
Hypothesized mechanism coordinating membrane phospholipid synthesis with cell growth in *Escherichia coli*. **A.** The role of PlsB in the *E. coli* phospholipid synthesis pathway. **B.** Our hypothesis predicts that during growth, the PlsB population is partitioned between an active form (either monomers or dimers) and inactive protein filaments. We propose that the degree of PlsB filamentation (and thus the level of PlsB activity and the rate of phospholipid synthesis) is determined by the density of phospholipids within the inner membrane: surplus phospholipids increase PlsB filamentation, while decreased phospholipid density (i.e. membrane stretching) decreases PlsB filamentation. This mechanism establishes an effective negative feedback loop that controls PlsB activity **(C)** and that maintains phospholipid density at a homeostatic level **(D)**.

As the first enzyme of the phospholipid synthesis pathway, PlsB activity determines the rate of membrane phospholipid synthesis. Thus, PlsB activity must be synchronized with growth. Phospholipid synthesis flux during steady-state growth is not controlled by concentrations of PlsB substrates (G3P and fatty acyl thioesters) or by PlsB concentration (5–8). Furthermore, PlsB overexpression does not increase phospholipid abundance (9, 10). Thus, PlsB activity must be regulated by an allosteric mechanism that activates and deactivates PlsB according to an allosteric signal. Despite extensive biochemical studies, potential allosteric mechanisms controlling PlsB activity have remained elusive (11–15). Intriguingly, an early study discovered that PlsB assembles into protein filaments that are readily detectable if PlsB is overexpressed (10). PlsB filaments consist of a repeating unit (likely PlsB dimers) helically arranged around a phospholipid core (16). While filamentous PlsB is enzymatically inactive, dissolution of PlsB filaments with a mild detergent restores activity. Whether filamentation coordinates PlsB activity with cell growth has remained unexplored.

Recently, filamentation is increasingly appreciated as a mechanism for regulating metabolic enzymes (17, 18). High-throughput imaging screens and biochemical experiments have found that many metabolic enzymes reversibly assemble into filaments (19, 20). Often, filamentation and inhibition are triggered by the accumulation of a product metabolite (21, 22). In these cases, filamentation enables negative feedback control over enzyme activity in response to product demand. Maintaining a fraction of the enzyme population in such an inactive form (described as “excess capacity” or “enzyme storage”) ensures that biosynthetic activity can be rapidly increased when needed (23).

How might filamentation play a role in regulating PlsB? By analogy to other filament-forming enzymes, PlsB might assemble into inactive filaments due to accumulation of excess membrane phospholipids. The fraction of cellular PlsB that is sequestered into inactive filaments might vary in proportion with the density of phospholipids within the inner membrane: a higher surface density of phospholipids would tend to increase the fraction of PlsB within a filament, decreasing overall PlsB activity, while decreased phospholipid content would liberate PlsB from filaments and increase the active fraction (**Fig. 1B**). This mechanism would establish a direct link between membrane phospholipid abundance and PlsB activity that acts as a classic negative feedback loop (**Fig. 1C**) and coordinates phospholipid synthesis with growth (**Fig. 1D**). Intriguingly, *in vitro* experiments suggest that excess phospholipids (generated by membrane compression) are extruded from membranes as small lipid filaments (24). Such lipid filaments might nucleate the formation of PlsB filaments, which have been found to contain an inner phospholipid core (25, 26)

Several lines of evidence are consistent with this model. As already stated, PlsB filaments are enzymatically inactive. Second, *E. coli* maintains a phospholipid surplus: *E. coli* protoplasts can be osmotically expanded before lysis occurs (27, 28), and *E. coli* continues to grow for a short time after phospholipid synthesis is inhibited (29). Third, PlsB is also present in excess: the maximum activity of PlsB far exceeds activity needed to provide *E. coli* with phospholipids (8). Finally, when phospholipid synthesis inhibition is relieved, the rate of phospholipid synthesis is transiently faster than the rate observed during steady-state growth, consistent with a brief period of uninhibited phospholipid synthesis in the absence of a phospholipid surplus (30).

However, direct evidence for the PlsB filamentation hypothesis is lacking. PlsB filaments have never been observed in wild-type cells. This is likely due to the comparatively low abundance of PlsB in *E. coli* (∼1000 PlsB/cell (31)). As protein filaments typically assemble in a cooperative manner (*i.e.* the energetic barrier for nucleating a new filament is higher than for extending an existing filament (32)), each cell might have at most 1-2 filaments. From these assumptions and estimates of PlsB filament geometry (26), a filament containing half the total PlsB population would be small (∼180 nm) and thus difficult to observe *in situ.* Here, we use fluorescent tagging to determine PlsB localization in wild-type cells. Our observations are consistent with our PlsB filamentation hypothesis: specifically, accumulation of membrane phospholipids triggers PlsB filamentation and inactivation, enabling negative feedback regulation of phospholipid synthesis and coordinating phospholipid synthesis with growth.

## Results

### PlsB localizes as punctate foci during growth

The ability of proteins to assemble into filaments *in vivo* has been identified in several studies by examining libraries of strains expressing fluorescently tagged proteins (33, 34). To enable PlsB localization, we fused a monomeric GFP variant (msGFP2) to the C-terminus of PlsB (35). As protein fusions may perturb localization and protein activity (36), we first investigated whether our fusion influenced PlsB activity. As PlsB is an essential enzyme, any detrimental effects of protein fusion on PlsB activity are expected to be reflected in the growth rate. Furthermore, localization artefacts may also arise when proteins are expressed above native concentrations. To ensure that PlsB-msGFP2 is expressed at native levels and that any localization observed is not a consequence of overexpression, we integrated the *plsB-msGFP2* gene into the chromosome of wild-type *E. coli* (NCM3722) at the *plsB* locus, replacing unlabelled PlsB while retaining transcriptional control by the *plsB* promoter. *E. coli plsB-msGFP2* grew normally in all tested media (**Fig. 2A**). To determine whether our fusion construct perturbs PlsB function or regulation, we quantified both membrane phospholipids and phospholipid intermediates using liquid chromatography coupled with mass spectroscopy (LCMS). The phospholipid composition of *plsB-msGFP2* cultures (determined by phospholipids/OD) and levels of phospholipid synthesis intermediates was identical to the parent NCM3722 strain in multiple conditions (**Fig. 2B**). Next, we quantified PlsB-msGFP2 using LCMS-based targeted proteomics. The abundance of PlsB peptides in *plsB-msGFP2* closely matched those measured in the parent NCM3722 strain, confirming that PlsB-msGFP2 is expressed from the chromosomal *plsB* locus at native levels (**Fig. 2C**). To confirm that msGFP2 labelling does not disrupt mechanisms that prevent overexpressed PlsB from increasing phospholipid content, we compared phospholipid abundance in our *plsB-msGFP2* strain in which we overexpressed PlsB-msGFP2 from a plasmid with NCM3722 overexpressing unmodified PlsB. While both PlsB and PlsB-msGFP2 overexpression slightly decreased total phospholipid abundance, the phospholipid content of the strain overexpressing PlsB-msGFP2 did not substantially differ from the strain overexpressing PlsB (**Fig. 2D**).

**Figure 2.**
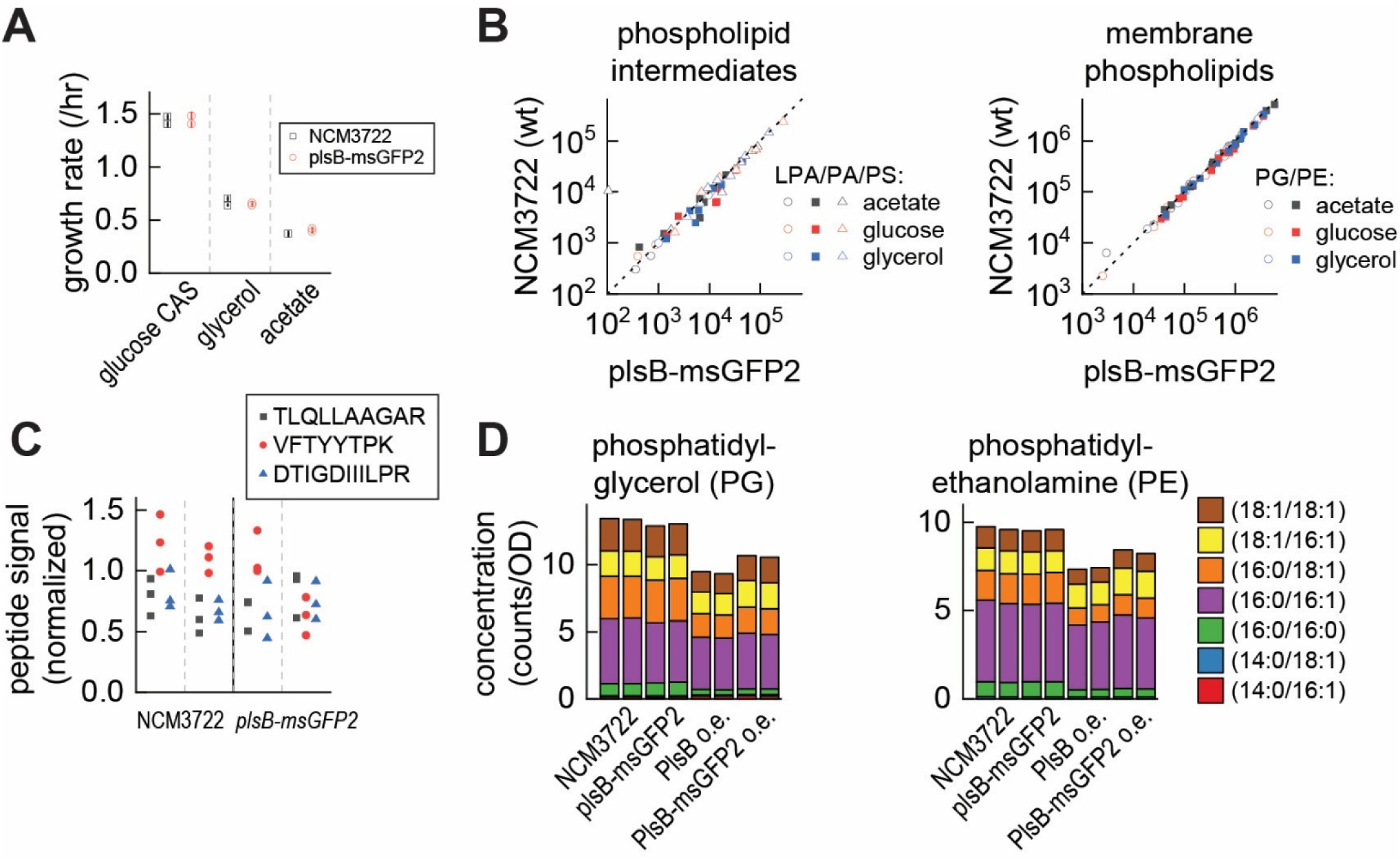
A msGFP2 fusion with PlsB retains activity. **A.** Growth rates of *E. coli* NCM3722 (wild-type) and NCM3722 *plsB-msGFP2* (PlsB-msGFP2 expressed from the chromosomal *plsB* operon) obtained in three defined media. Two biological replicates of each strain in each condition are shown. **B.** Abundance of phospholipid synthesis intermediates (LPA, PA, and PS) and membrane phospholipids (PG and PE) are unaffected by expression of PlsB-msGFP2. Data obtained from one biological replicate in each condition. **C.** Cellular concentration of PlsB-msGFP2 is equivalent to concentration of PlsB in NCM3722. Triplicate data obtained using LCMS-based peptide quantification from two biological replicates of each strain are shown. **D.** Overexpression of PlsB in NCM3722 and PlsB-msGFP2 in *plsB-msGFP2* generate similar membrane phospholipid profiles. Duplicate data from one biological replicate of each condition is depicted.

To determine how PlsB localizes during steady-state growth, we imaged exponential phase *plsB-msGFP2* cells from a defined rich medium culture (MOPS / 0.2% glucose / 0.1% CAS amino acids). Cells were transferred to an agar pad supplemented with the same medium and imaged using live-cell epifluorescence microscopy. Fluorescent foci were found within 95% of the population, with diffuse fluorescence appearing within the cytoplasm (**Fig. 3A&B**). Most PlsB foci localized at the cell poles (**Fig. 3C**). PlsB foci were present at comparable levels in all growth medium tested, indicating that foci are not an artefact of a specific condition. To determine whether foci localization is an artefact caused by the msGFP2 fluorescent tag, we also imaged PlsB localization using a C-terminal HaloTag fusion. PlsB-HaloTag also localized as punctate foci at a similar frequency as PlsB-msGFP2 (**Fig. 3D**). Finally, we exchanged msGFP2 with the HA epitope and imaged PlsB using immunofluorescence. PlsB-HA was expressed from a plasmid in wild-type NCM3722 cells that were subsequently fixed and imaged with anti-HA antibodies. Overexpressed PlsB-HA also localized as punctate foci in wild-type NCM3722 (**Fig. 3E**). Thus, PlsB foci are not an artefact of specific tags or overexpression.

**Figure 3.**
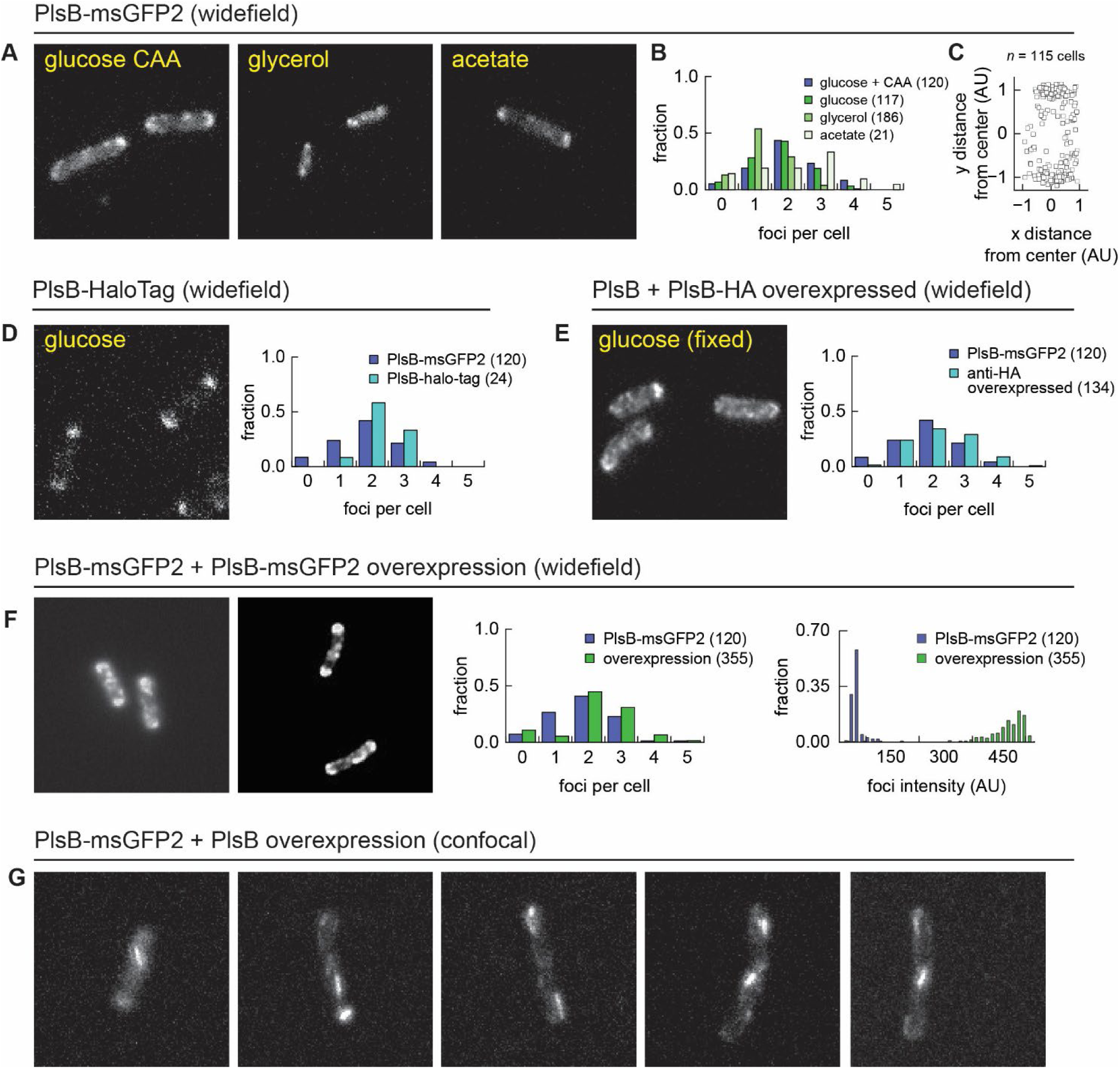
PlsB localizes as foci during steady-state growth. **A.** Live-cell fluorescent microscopy images of *E. coli* NCM3722 *plsB-msGFP2* on agar pads supplemented with defined medium. **B.** PlsB foci form at similar frequencies across conditions and usually localize at cell poles **(C)**. **D.** PlsB foci are also observed with different protein fusions, and in fixed cells **(E)**. **F.** Plasmid-based overexpression of PlsB-msGFP2 does not affect the number of PlsB foci detected per cell but increases the brightness of the foci. **G.** Overexpression of unlabelled PlsB in *plsB-msGFP2* background allows resolution of fluorescent structures resembling filaments. For all panels, the numbers of analysed cells are provided in the figure legends.

### Fluorescence intensity of PlsB foci increases with PlsB expression

The observation that fluorescently tagged proteins localize as foci is consistent with assembly into filaments or other structures such as chemotaxis receptor arrays (37). In many cases, the degree of oligomerization is dependent upon protein concentration, with increased oligomerization correlating with increased protein concentrations. In contrast, if the number or size of foci do not correlate with protein expression, any foci detected may not indicate protein oligomers. For instance, the primary phospholipid synthesis enzyme in *Bacillus subtilis,* PlsX, localizes as foci in some conditions, but foci number did not increase in parallel with PlsX expression (38). To determine whether the PlsB foci represent protein oligomers, we expressed the PlsB-msGFP fusion from a plasmid-borne inducible promoter in our *plsB-msGFP2* strain. Inducing expression of PlsB-msGFP2 did not increase the number of fluorescent foci observed but dramatically increased the foci brightness (**Fig. 3F**), consistent with PlsB foci indicating formation of an oligomeric complex. Thus, the effects of increased PlsB-msGFP2 expression are also consistent with assembly of multimeric complexes.

### C-terminal tagging may perturb formation of long filament structures

The foci observed following overexpression of PlsB-msGFP2 did not appear as elongated filaments as expected from previous studies (**Fig. 3F**). To determine whether the GFP fusion affects the formation of filaments, we overexpressed unlabeled PlsB in the *plsB-msGFP2* strain and imaged our cells using confocal microscopy. In contrast to the foci detected within cells overexpressing PlsB-msGFP2 alone, we observed linear fluorescent structures more consistent with filaments (**Fig. 3G**). This implies that the presence of a C-terminal tag may adversely affect either the assembly of PlsB into an orderly filamentous array or the parallel packing of multiple PlsB filaments. However, immunofluorescence imaging of overexpressed PlsB-HA also appear as foci. Thus, modification of PlsB on the C-terminus may affect the assembly of PlsB filaments.

### Inhibition of fatty acid synthesis delocalizes PlsB

Despite the unclear effect of C-terminal tagging on the formation of orderly PlsB filaments, the consistent foci-forming behaviour of PlsB is consistent with assembly into some oligomeric structure. However, the functional relevance of the foci is unclear. Fluorescent tagging often introduces artefacts that do not reflect the native location of untagged proteins (39, 40). One approach for testing connections between foci formation and enzyme function is to determine whether foci formation is responsive to metabolic perturbations linked with the function of the enzyme (41). Specifically, if the PlsB foci we observe reflect oligomerization in response to phospholipid membrane abundance, perturbations to membrane abundance or phospholipid metabolism are expected to affect localization. Conversely, if the PlsB foci instead are the consequence of a labelling artefact (e.g. fusion-induced aggregation), the PlsB foci are expected to be unresponsive to perturbations in membrane metabolism.

We tested whether PlsB localization is relevant to its physiological function by inhibiting fatty acid synthesis. Triclosan is a fatty acid synthesis inhibitor that acts on the 2*E*-enoyl-acyl-ACP reductase FabI (42). Triclosan was added to cells growing in liquid culture, which were subsequently transferred to an agar pad also containing growth medium and triclosan for live-cell imaging. Triclosan greatly reduced the number of PlsB foci and delocalized fluorescence throughout the cytoplasm (**Fig. 4A**). A similar result was obtained using the fatty acid synthesis inhibitor cerulenin, which inactivates the β-keto-acyl-ACP synthases FabB and FabF (43). To determine whether PlsB foci respond to milder perturbations of fatty acid synthesis, we overexpressed the *E. coli* thioesterase TesA without its periplasmic trafficking signal. When localized to the cytoplasm, TesA hydrolyses long-chain fatty acid thioesters (including acyl-ACP PlsB substrates) (44) and reduces the abundance of membrane phospholipids (45) while allowing growth to continue. Overexpression of TesA in *plsB-msGFP2* dispersed PlsB-msGFP2 and reduced the foci count (**Fig. 4A**). To determine whether PlsB foci dispersal is caused by generic cell growth arrest rather than a specific consequence of fatty acid synthesis inhibition, we imaged *plsB-msGFP2* after adding protein translation inhibitors chloramphenicol and mupirocin. Cells treated with either translation inhibitor ceased to grow but retained PlsB foci (**Fig. 4B**), indicating that PlsB delocalization is not a generic response to growth arrest but is a consequence of fatty acid synthesis inhibition. Thus, PlsB consistently delocalizes in response to perturbations in fatty acid and phospholipid synthesis.

**Figure 4.**
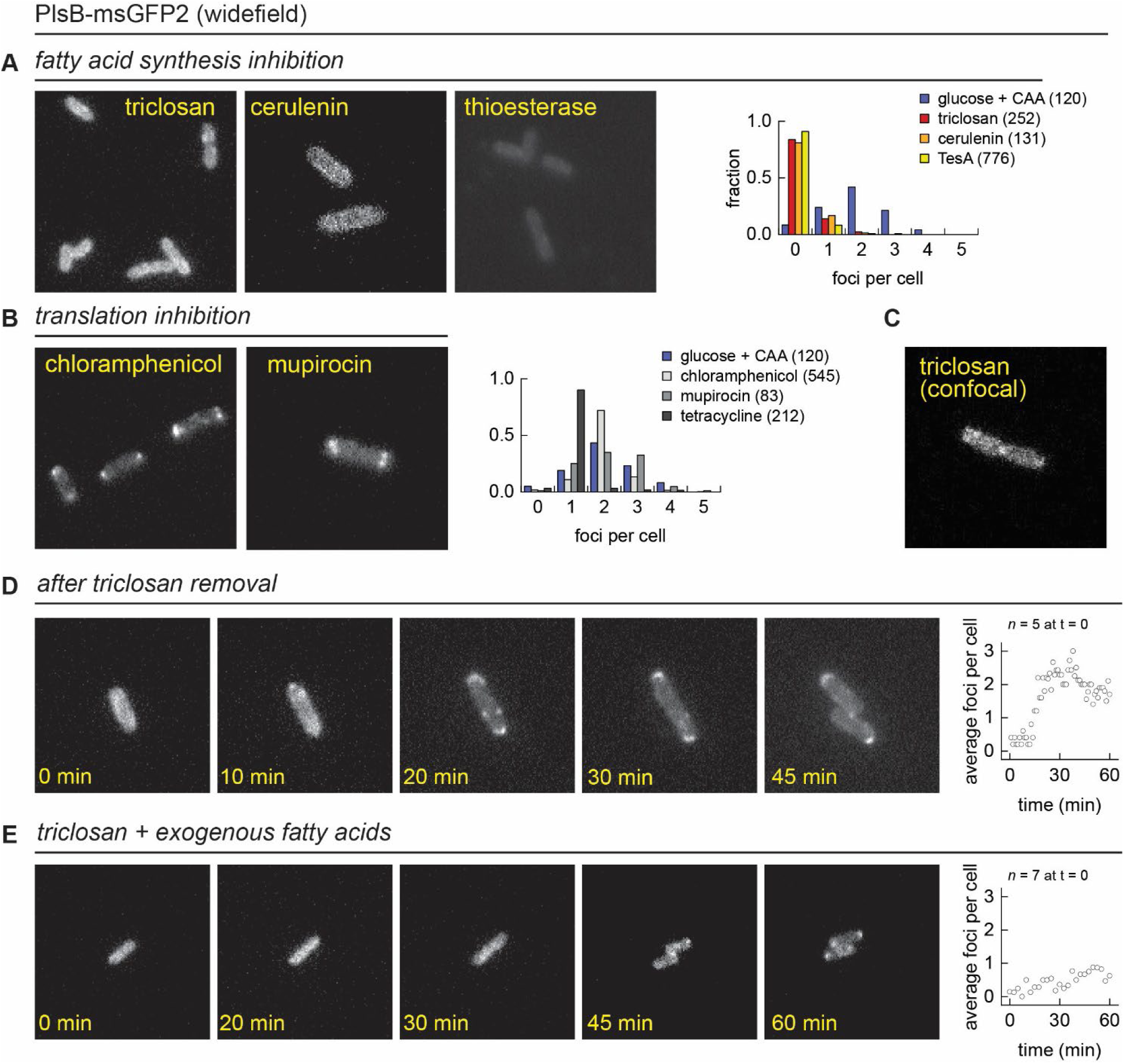
PlsB foci respond dynamically to phospholipid abundance. **A.** PlsB-msGFP2 delocalizes into the cytoplasm after membrane synthesis inhibition. Images of *plsB-msGFP2* cells grown in MOPS/0.2% glucose 0.1% CAS amino acids treated with triclosan, cerulenin, or expressing the thioesterase TesA and transferred to agar pads supplemented with identical medium and antibiotics. **B.** PlsB-msGFP2 foci do not substantially respond to translation inhibitors, indicating that delocalization is not a generic response to growth arrest. **C.** A confocal image of triclosan-treated *plsB-msGFP* suggests that PlsB-msGFP2 delocalizes into the cytoplasm and does not associate with the membrane. **D&E.** PlsB-msGFP2 foci reappear when membrane synthesis is restored by either removing triclosan (**D**) or by adding free fatty acids palmitate and *cis-*vaccenate (**E**). For all panels, the numbers of analysed cells are provided in the figure legends.

The cytoplasmic localization of PlsB observed when phospholipid synthesis is inhibited is unexpected. While PlsB is a peripheral membrane protein and not integrated into the membrane, it is generally regarded as associating stably with the membrane. To further investigate PlsB localization with higher spatial resolution, we imaged triclosan-treated *plsB-msGFP2* using confocal microscopy. Consistent with observations from our wide-field images, PlsB-msGFP2 in triclosan-treated cells is also cytoplasmic and not membrane-associated (**Fig. 4C**). We note that the phosphatidylserine synthesis enzyme PssA is also a peripheral membrane enzyme that reversibly associates with the membrane in response to electrostatic charge and substrate availability (46).

### PlsB foci gradually reform after phospholipid synthesis is restored

We next tested whether resumption of fatty acid synthesis might cause PlsB foci to reform. Triclosan-treated cells in liquid culture were pelleted, resuspended in fresh medium, and plated on agar containing growth medium and imaged over time. Growth resumed immediately upon triclosan removal, while PlsB remained dispersed at the beginning of imaging. However, after ∼30 minutes of growth, foci reappeared, coincident with the time of first cell division (**Fig. 4D**). Foci reappearance is highly synchronized across the population. Thus, both formation and dispersion of PlsB are reversible and responsive to phospholipid metabolism.

To further investigate PlsB foci formation, we restarted phospholipid synthesis in triclosan-inhibited cells by feeding exogenous fatty acids palmitate and *cis-*vaccenate. Exogenous fatty acids are converted by the fatty acid-CoA ligase FadD to acyl-CoA, which are substrates for phospholipid synthesis by PlsB and PlsC. Fatty acid-fed cells resumed growth at a much slower rate; nevertheless, PlsB foci steadily accumulated in the population, although less synchronized than when phospholipid synthesis is restarted by triclosan removal (**Fig. 4E**). We speculate that the slower growth rate and delayed appearance of foci is due to the inability to synthesize lipopolysaccharides, which require lipid precursors that cannot be supplied or obtained from exogenous fatty acids. Thus, PlsB consistently localizes as foci when membrane synthesis is unperturbed during steady-state growth but disperses when phospholipid abundance is reduced.

## Discussion

Indirect evidence from previous studies is consistent with our hypothesis that PlsB activity (and thus phospholipid synthesis) is regulated by reversible filamentation. First, PlsB concentrations in wild-type cells are far above levels required to provide sufficient phospholipids for steady-state growth (1, 8, 47); second, overexpressed PlsB forms enzymatically inactive filaments (25, 26); finally, many metabolic proteins assemble into enzymatically inactive filaments in the presence of their products (32, 48). Our observations from live-cell imaging experiments support our hypothesis: PlsB localizes as a combination of foci and distributed fluorescence, consistent with the formation of a non-diffusive, membrane-associated state such as a filament. Furthermore, PlsB foci intensity varies in proportion with PlsB expression, consistent with increased PlsB oligomerization, and contrasts with results obtained from fluorescently labelled PlsX in *B. subtilis* cells (38). However, while fluorescent foci are consistent with our hypothesis, many other mechanisms (natural and artefactual) also cause labelled proteins to appear as foci (36). Thus, our strongest evidence is that PlsB localization responds reversibly to membrane abundance in a manner predicted by our hypothesis. Specifically, PlsB foci disappear when membrane phospholipids are depleted and reappear when phospholipid limitation is alleviated.

While our live-cell microscopy data support our hypothesis, some questions remain. First, the foci that we observe may not necessarily indicate the formation of PlsB filaments. For instance, the foci may simply reflect increased localization of PlsB monomers or dimers rather than an oligomeric structure. This may be due to a preference for binding in regions of anionic phospholipids or for the curved membrane at the cell poles. Issues also arise from our use of protein fusions to localize PlsB in live cells. C-terminal fusions appear to compromise filament formation. This may be caused by the C-terminal extensions disrupting contacts or tweaking the alignment of monomers and preventing helical packing observed in untagged PlsB. Alternatively, C-terminal tagging may disrupt the arrangement of multiple PlsB filaments into parallel bundles. Unfortunately, solutions such as appending tags to the N-terminus, or inserting a tag in an unstructured loop (an approach successfully used to study MreB localization (49)) are not straightforward to apply to PlsB: the N-terminus lies at the interface with the membrane, and the Alphafold-predicted structure of PlsB does not reveal any obvious unstructured loops that might allow the insertion of a bulky fluorescent label (although a smaller label may be tolerated). Interestingly, although formation of orderly filaments appears to be disrupted, C-terminal tagging does not dramatically affect PlsB function or regulation: growth rate and phospholipid abundance of *plsB-msGFP2* are unaffected. This may even be interpreted as an indication that PlsB and phospholipid homeostasis do *not* in fact rely upon PlsB filamentation. However, multiple studies that have established the role of filamentation in regulation of other metabolic enzymes found that disabling filamentation caused surprisingly subtle effects upon fitness: for instance, disrupting glucokinase filamentation lowered fitness only during growth transitions, not during steady-state growth (41).

To directly test whether filamentation is required for membrane synthesis regulation, PlsB mutants that are unable to form filaments should be created and tested *in vivo*. Unfortunately, a single helical turn of a PlsB filament requires more PlsB monomers (25, 26) than current oligomer structure prediction tools allow (50). Thus, PlsB filament structures must be experimentally determined at sufficient resolution to identify residues that stabilize filaments. Expression of filamentation-deficient PlsB mutants would be expected to increase the phospholipid/protein ratio well above wild-type levels would provide additional direct support for our hypothesis. Furthermore, experiments with lipid bilayers supported by a flexible surface could determine whether PlsB filaments are nucleated by extruded lipid filaments, an approach used to study membrane remodelling by BAR proteins (51).

## Methods

### Strains and growth conditions

*Escherichia coli* K-12 strain NCM3722 (CGSC #12355) was used for all experiments. Plasmids and primers used are listed in Tables 1 and 2, respectively, while the amino acid sequences of protein fusions created are in Table 3. *plsB-msGFP2* was cloned into BglBrick plasmid pBbE5k, sub-cloned into pDHL940 to fuse with a kanamycin resistance gene, and subsequently inserted together with the resistance cassette into the NCM3722 chromosome by lambda-red recombination (52). The kanamycin resistance cassette was subsequently removed. The *plsB-HaloTag* strain was constructed in a similar manner. Cells were cultured on MOPS minimal media (53) with 0.2% w/v carbon source (glucose, glycerol, or acetate) or with 0.2% glucose and 0.1% CAS amino acids. Growth was monitored with optical density measurements (Ultrospec 10 Cell Density Meter, GE Healthcare). For microscopy experiments, 5 mL cultures were incubated at 37°C in while rotating at 400 rpm. For cultures prepared for experiments requiring larger volumes (growth rate measurements and LCMS analysis), cells were grown in Erlenmeyer flasks containing a magnetic stir bar. Culture flasks were partially submerged in in a Grant Instruments Sub Aqua Pro water bath maintained at 37 °C and stirred (1200 rpm) with submerged 2mag MIXdrive 1 Eco and MIXcontrol 20 stir plates. PlsB expression plasmids (PlsB, PlsB-msGFP2, and PlsB-HA) were constructed by inserting the PlsB or PlsB fusion gene into a pBbE5k backbone (54).

**Table 1.** Plasmids in this study.

| Plasmid | Role | Source |
| --- | --- | --- |
| pQE-60NA-msGFP2 | msGFP2 sequence | AddGene (plasmid #135301) |
| pDHL940 | HaloTag sequence and antibiotic cassette for recombination | Mehmet Berkmen (57) |
| pBbE5k | BglBrick vector (lacUV5 promoter, ColE1 origin, kanamycin resistance marker) | AddGene (plasmid #35279) |
| pBbA2a | BglBrick vector (pTet promoter, p15a origin, ampicillin resistance marker) | AddGene (plasmid #35325) |
| pBbE5k-PlsB-msGFP2 | PlsB-msGFP2 overexpression | This study |
| pBbA5a-TesA | Cytoplasmic thioesterase expression | This study |
| pBbE5k-PlsB | PlsB overexpression | This study |
| pBbE5k-PlsB-HA | HA-tagged PlsB overexpression | This study |

**Table 2.**
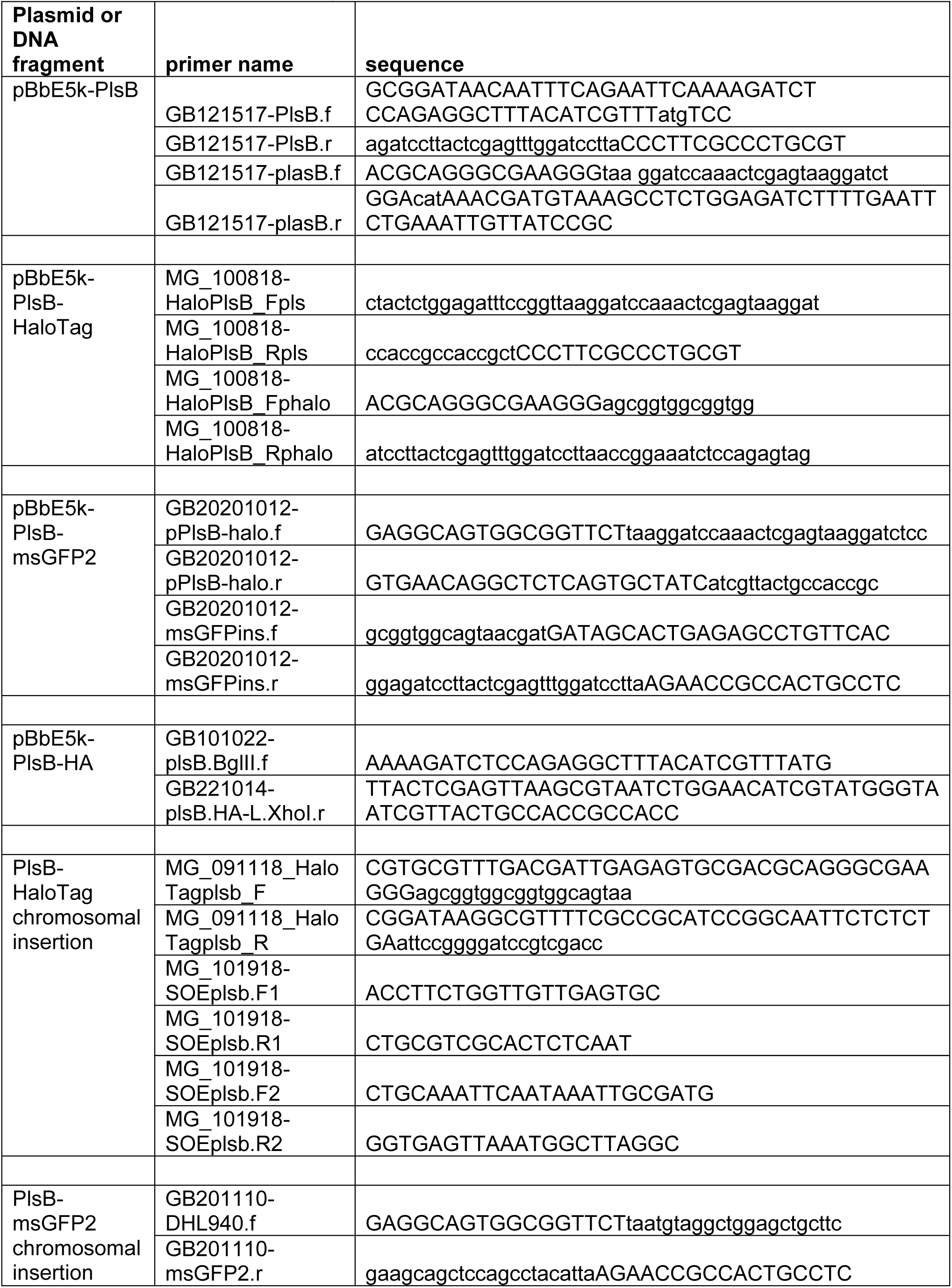
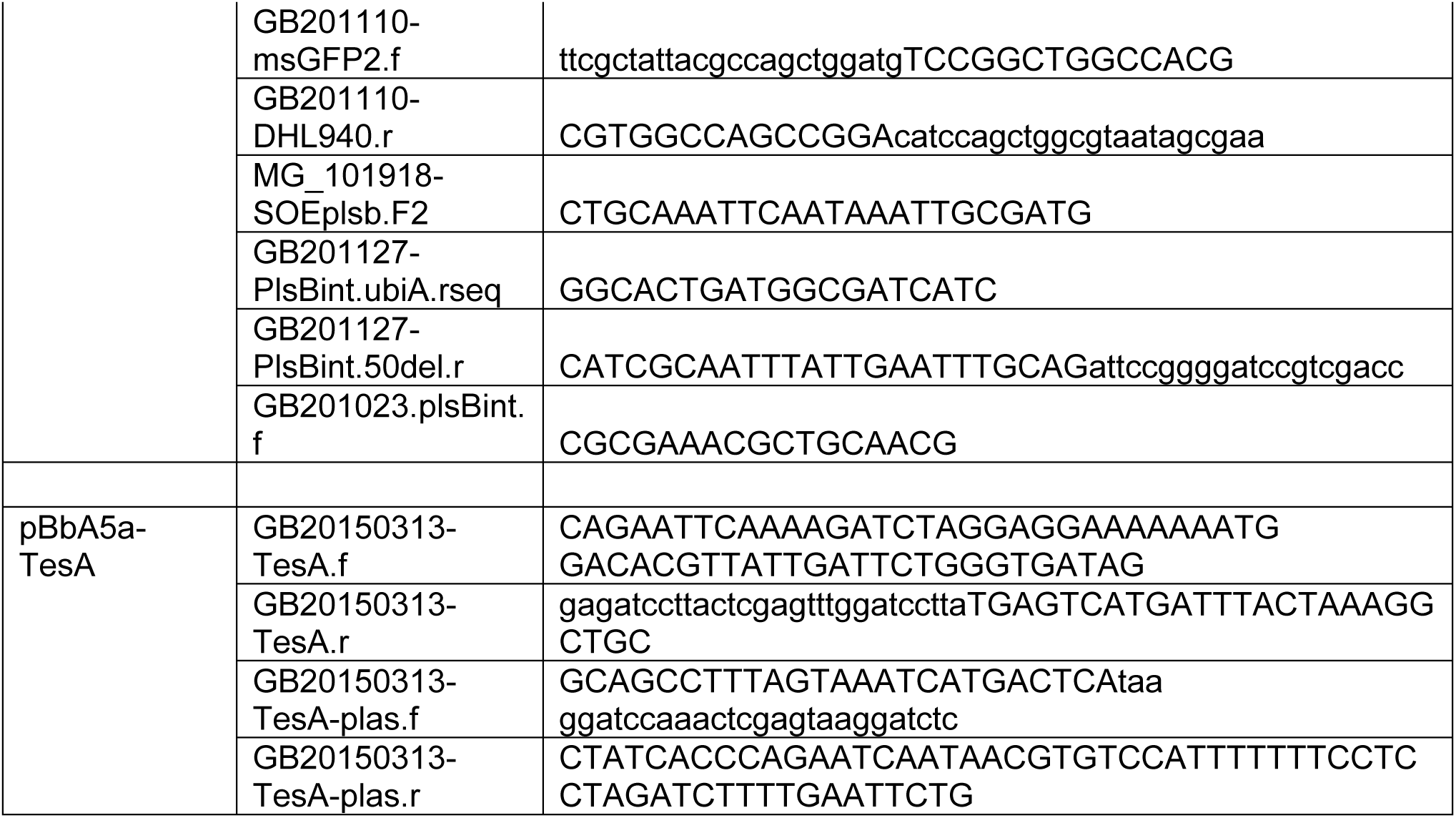
Primers used in this study.

**Table 3.**
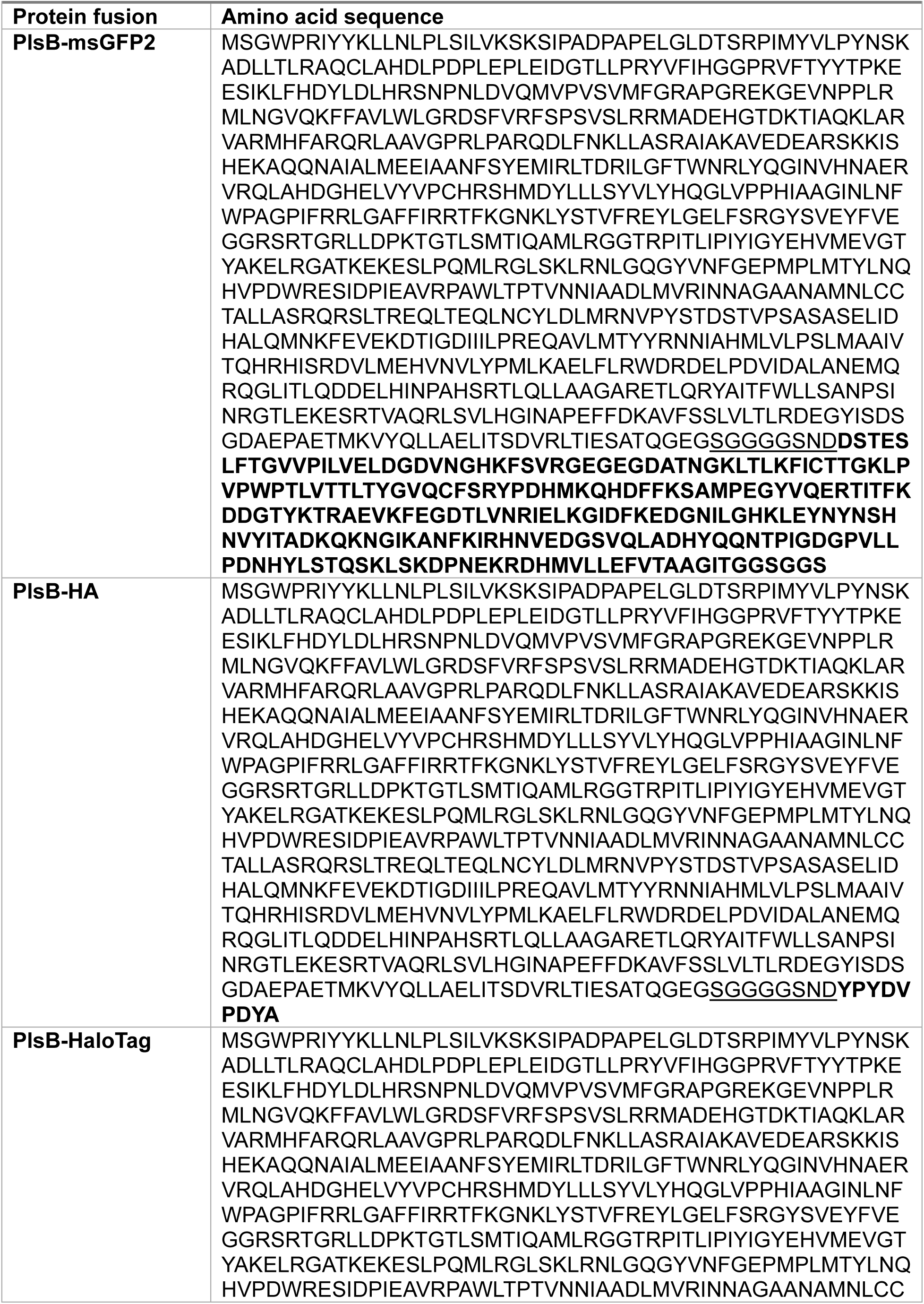
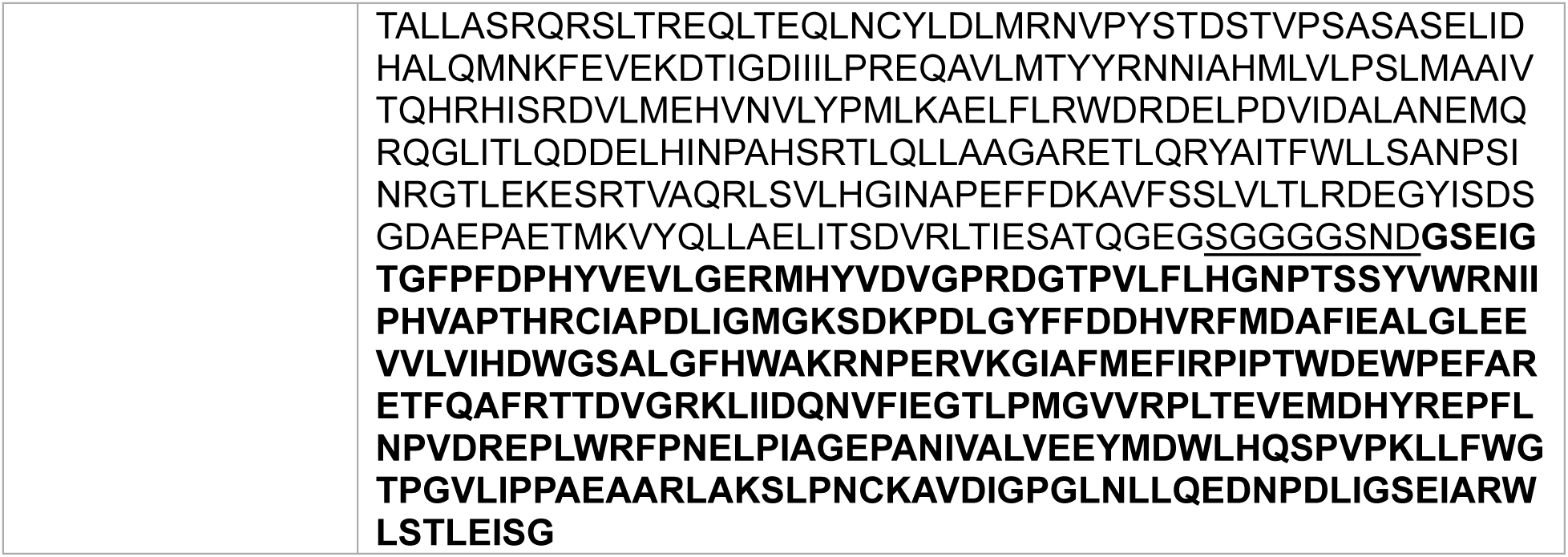
Amino acid sequences of PlsB fusions. Linker sequences are underlined while fusion tags are in bold text.

### Culture sampling and LCMS analysis

1 mL samples were rapidly removed from exponential-phase cultures and quenched by pipetting immediately into ice-cooled tubes containing 0.25 mL 10% trichloroacetic acid. Quenched cells were pelleted and stored at -80 °C until analysis. LCMS methods used for phospholipid and PlsB quantification are fully described in reference (8).

### Live-cell fluorescence microscopy

The cultures used for microscopy were prepared at outline in the “culture conditions” section. To ensure the bacteria grow during the imaging they were placed on agar pads. To make these pads, first a larger batch of agarose was prepared using 1.5 g low gelling point agarose dissolved in 50 mL milli-Q water for a final concentration of 3%. This agar was then autoclaved to ensure sterility. Afterwards, the full volume was divided into 0.5 mL aliquots which were stored at -20°C. For each experiment, one of these aliquots was used and remelted at 80 °C in a heating block. Once melted, 0.5 mL of a MOPS medium prepared at 2x concentration is added. The solution is mixed through vortexing and briefly (less than 5 min) placed back in the heating block to ensure it does not gel in the tube. Three 15 x 15 mm glass coverslips are place on a flat piece of parafilm and 0.3 mL of agar is pipetted on each of these. A second coverslip is then gently placed on top of the agar droplet, spreading it out to a (near) uniform shape. These are covered and left to gel, after they have solidified they are placed at 37 °C to pre-heat.

Microscopy slides were prepared in two different ways, depending on the microscope used. For widefield fluorescence microscopy, a prepared and pre-heated agar pad is taken and one of the coverslips peeled off. A small amount of culture (1 – 2 µL) is then placed onto the agar pad, which is then placed agar side down onto a larger 22 x 50 mm coverslip. If used for a timelapse, the sides are sealed to prevent drying of the agar pad during the acquisition. For confocal microscopy there is a slight variation on the above protocol, instead of placing the agar onto a 22 x 50 mm coverslip, they are instead placed onto a microscopy dish with a lid and a small amount of wet tissue paper inside to prevent dessication.

Widefield microscopy was done initially on an Inverted Olympus IX81 equipped with a Andor Luca R camera (EM-CCD, 104×1002 pixel chip size, 8×8µm pixel size, 14-bit dynamic range, 65% quantum efficiency at 600nm), CoolLED pE-4000 fluorescence excitation source, 100×1.45 NA oil objective (0.13 mm WB), GFP filter cube (ET-EGFP, 470/40 excitation filter, T515lp dichroic filter cut-off, 525/50 emission filter, reference: C229863), phase ring and heating box. The microscope was heated to 37°C before the start of the experiment, at least 30 minutes in advance to ensure a stable temperature. At the start of acquisition the 10x or 20x objective was used to check alignment of the phase ring and make adjustments if needed. Positions were chosen keeping in mind a sufficient number of cells in the field of view and no or minimal non-cell debris. Images were taken in two channels, one in phase contrast (50 ms exposure) and one for GFP (1000 ms exposure, 490 nm, 40% intensity). Phase contrast imaged were taken through the GFP filter cube to save time switching blocks. Where relevant, Z-stacks were taken using a 0.2 µm interval between slices. For timelapses, multiple positions are marked and imaged at 2.5 minute intervals, generally for at least 4 hours but up to 8 hours.

Confocal microscopy is done on a Nikon Eclipse Ti microscope with a 488 nm laser (confocal-CW, 16 mW), a 100x SR Apo TIRF 1.49 NA oil objective (0.12 mm WB), a 482/35 fluorescence emission filter, a EM-CCD Andor iXon X3 DU897 detection device (512×512 pixel chip size, 16×16 µm pixel size, 14 bit dynamic range, >90% quantum efficiency at 600nm), and Tokai Hit sample incubator chamber. The sample is incubated at 37°C during imaging. Suitable locations are found by looking at the sample using the Resonant scanning. Once a position is found a high resolution image is taken using Galvano scanning.

### HaloTag imaging

Cultures of *plsB-HaloTag* grown to OD600 of ∼0.6, incubated with 250 nM JF635 dye for 1 hour at 37 °C, and washed three times with medium before imaging.

### Immunofluorescence

An NCM3722 pBbE5K-PlsB-HA culture was induced with 100 µM IPTG and grown for approximately 1 hour until OD600 of 0.3 – 0.4. Following a gentle fixation protocol (55), the cells were subsequently fixed for 15 minutes with formaldehyde and glutaraldehyde (final concentrations 2.8% and 0.04%, respectively), washed twice with PBS and concentrated 100-fold by centrifugation and resuspension in PBS. The dense cell suspension was incubated for 45 min with 1% Triton X-100 and washed three times with 150 µL PBS. Next, the cells are incubated 45 min in 150 µL PBS containing 100 µg/mL lysozyme and 5 mM EDTA, followed by three washes with 150 µL PBS and stored at -20°C until ready for labelling. For antibody labelling, cells were incubated for 1 hour at 37 °C with 0.5 mg/mL antibody (anti-HA rabbit recombinant monoclonal antibody labeled with Alexa Fluor 488, Abcam EPR22919-101). Cells are washed three times with 150 µL PBS containing 0.05% Tween-20, resuspended in 150 µL PBS and analyzed using fluorescence microscopy.

### Image analysis

To augment the qualitative analysis of microscopy images we developed an image analysis pipeline. The primary goal of this pipeline is to count the number of foci within each cell. This requires detection of individual cells and the subsequent detection of the foci within each cell. The first step is to create image stacks of both the phase contrast and fluorescent images separately. The phase contrast images are loaded into Fiji. The MicrobeJ plugin (56) is then used to segment the cells. The first segmentation is automatic, and this is then followed by manual refinement where necessary. Manual refinement is almost exclusively reserved for eliminating false positives or for separating touching cells. Cells touching the edge are discarded and for all remaining cells the coordinates of their contours are exported. Next, both these contours and the fluorescent images stack are loaded into a custom-made Matlab script. Here the contours are used to find each individual cell in the fluorescent images. All other pixels not defined as cell are considered background and the average background value is subtracted across the image. Next, each cell is analyzed individually. First the cytoplasmic background is estimated by taking the median intensity of cell. This value is then multiplied with a set factor (1.7) to create a threshold. All pixels higher than this threshold are located and further filtered on cluster size. Only clusters of more than 5 pixels are considered a foci. From this we can then collect the number of foci per cell, the number of pixels (i.e. size) and the total intensity of the foci. Next, the centroid of the cell is found as well as the centroid of each foci to determine where in the cell they are located. Besides the exact location of each foci, they are also classified as polar or non-polar. For this determination, foci are considered to be located at the poles when located at the outer 10% of the cell length on either end. Finally, the cell length and width are also measured. All this data is then collected and exported for further analysis.

## Acknowledgements

We thank Mehmet Berkmen for sharing HaloTag plasmid pDHL940, Carlas Smith and Michal Shemesh for expertise and assistance with microscopy, Jeremie Capoulade for performing pilot experiments, and Tanneke den Blaauwen and Jolanda Verheul for assistance with cell fixation and immunofluorescence. We thank Ariane Briegel and Arjen Jakobi for encouragement and insightful suggestions, and the entire Bokinsky Lab for stimulating discussions and support.

## Author Contributions

JB: designed and performed experiments, built image analysis pipeline, evaluated data. DF: collected samples for LCMS experiments. MG: constructed *plsB-HaloTag* strain and performed HaloTag imaging experiments. AS: performed initial experiments confirming TesA-driven PlsB delocalization. AZ-D: LCMS analysis. GB: supervised project, constructed *plsB-msGFP2* strain, evaluated data. JB and GB wrote the manuscript. All authors approved the final manuscript.

## Notes

### Competing Interest Statement

The authors have declared no competing interest.

### Summary of Updates

Minor revisions for clarity. Primer sequences used to construct plasmids and DNA constructs for recombination added.

